# IRE1α drives lung epithelial progenitor dysfunction to establish a niche for pulmonary fibrosis

**DOI:** 10.1101/2021.09.16.460705

**Authors:** Vincent C. Auyeung, Michael S. Downey, Jaymin Kathiriya, Maike Thamsen, Talia A. Wenger, Bradley J. Backes, Alexis Brumwell, Harold A. Chapman, Dean Sheppard, Feroz R. Papa

## Abstract

Idiopathic pulmonary fibrosis (IPF) is a disease of progressive interstitial fibrosis, which leads to severe debilitation, respiratory failure, and death. In IPF, environmental exposures interact with genetic risk factors to engender critical patho-etiological events in lung epithelial cells, including endoplasmic reticulum (ER) stress and TGFβ signaling, but the interactions between these disparate pathways are not well understood. We previously showed that kinase inhibitors of the IRE1α bifunctional kinase/RNase—a central mediator of the unfolded protein response (UPR) to ER stress—protected mice from bleomycin-induced pulmonary fibrosis. Here we show that a nanomolar-potent, mono-selective kinase inhibitor of IRE1α (KIRA8) decreases ER-stress induced TGFβ signaling and the senescence-associated secretory phenotype (SASP) in the lung epithelium after bleomycin exposure. A recently-described subset of “damage-associated transient progenitors” (DATPs) display IRE1α-regulated pathological gene signatures that are quelled by KIRA8, *in vivo*. After injury, these cells uniquely express integrin αvβ6, a key activator of TGFβ in pulmonary fibrosis. KIRA8 inhibition of IRE1α decreases both DATP number and *Itgb6* expression in remaining cells, with a decrease in local collagen accumulation. Single-cell RNA sequencing from IPF lungs revealed an analogous *Itgb6+* cell population that may also be regulated by IRE1α. These findings suggest that lung epithelial progenitor cells sit at the center of the fibrotic niche, and IRE1α signaling locks them into a dysfunctional state that establishes and perpetuates pathological fibrosis.

## Main Text

Idiopathic pulmonary fibrosis (IPF) is a disease of progressive interstitial fibrosis, which leads to severe debilitation and eventually respiratory failure and death. An emerging paradigm is that environmental exposures interact with genetic risk factors to engender multiple events in lung epithelial cells, including endoplasmic reticulum (ER) stress and the unfolded protein response (UPR), premature telomere shortening, and senescence. These ultimately trigger pro-fibrotic signals, including integrin αvβ6 and TGFβ, Wnts, and PDGFα.(1)

An intracellular signaling pathway—the UPR—becomes activated under ER stress, and classically induces components of the ER protein folding machinery to adaptively restore protein folding homeostasis. But under conditions of severe, irremediable ER stress, the UPR actively promotes a continuum of maladaptive cell fate outcomes, including trans-differentiation, senescence, and apoptosis.(2) These cell fate switches are controlled in part by the ER stress sensor IRE1α, an ER transmembrane protein containing an ER luminal domain that is responsive to protein-folding status in the ER. On its cytosolic face, IRE1α features interlinked kinase and RNase domains that transduce the ER stress signals of immature/unfolded protein buildup in the ER to the cytosol and nucleus, where corrective programs are launched. Upon sensing ER stress, IRE1α kinase domains self-associate and *trans*-autophosphorylate, which activates the RNase domain to initiate frameshift splicing of the mRNA encoding the adaptive XBP1 transcription factor through site-specific endonucleolytic cleavage. However, continuous or high-level ER stress causes IRE1α kinase/RNAse domains to also degrade a plethora of mRNAs localizing to the ER membrane and some select micro-RNAs in a process dubbed “RIDD”— Regulated IRE1α-Dependent Decay—leading to the aforementioned maladaptive cell fate outcomes.(3–5)

ER stress and consequent UPR activation occurs in familial forms of IPF in which mutations in surfactant protein cause its misfolding, or protein trafficking is perturbed by loss-of-function mutations in the BLOC-3 complex, leading to injury to lung epithelial cells.(6, 7) In mice, ER stress can be induced in alveolar epithelial cells by knockout of a key chaperone, BiP, or knock-in of a surfactant protein C mutation (C121G) found in human IPF patients; both are sufficient to induce lung injury and fibrosis.(8, 9) Activation of the UPR has also been described in non-familial IPF, where the cause of UPR activation appears to be more complex and includes viral infection, DNA damage, senescence, and localized hypoxia.(10–12)

IRE1α is of particular interest in the pathophysiology of IPF because DNA damage, senescence, and pathogen-associated molecular patterns have each been shown to trigger selective IRE1α activation, with little or no activation of other arms of the UPR mediated by PERK or ATF6.(13–16) Even more remarkably, senescence and DNA damage appear to induce RIDD, an activity associated with hyperactivated IRE1α.(15, 16) RIDD mediates a range of maladaptive cellular outcomes that have been implicated in pulmonary fibrosis, ranging from TGFβ induction and inflammatory cytokine production to senescence and apoptosis.(5, 17–20) These observations suggest that IRE1α may be a key player in a network of mutually reinforcing pathways that conspire to promote pathological fibrosis.

Since upon autophosphorylation IRE1α’s kinase domain rheostatically activates its RNase domain, we previously developed small molecule ATP-competitive kinase inhibitors to allosterically attenuate the RNase catalytic activity, called *<u>K</u>*inase *I*nhibiting *<u>R</u>*Nase *<u>A</u>*ttenuators—”KIRA”s.(21) We showed that KIRAs could protect mice from experimental pulmonary fibrosis induced by bleomycin exposure, even when given as late as two weeks after bleomycin exposure.(22) To decipher the basis of this therapeutic efficacy, in this study we used ribosome-tag sequencing to better understand how IRE1α is wired into profibrotic gene expression programs in epithelial cells after bleomycin exposure. We found that a nanomolar-potent, mono-selective kinase inhibitor of IRE1α—KIRA8—decreased TGFβ signaling and the senescence-associated secretory phenotype (SASP), while increasing levels of the microRNA miR-17, which we had previously shown to be capable of dampening a maladaptive UPR.(5) Analysis of single-cell transcriptomes revealed that these maladaptive UPR programs are concentrated in a discrete population of damage-associated transient progenitor cells (DATPs) that can serve as a pool of progenitor cells to repair the lung, but are instead stalled in a transitional state after injury and paradoxically drive fibrosis. IRE1α regulated both the number and dysfunctional phenotype of DATPs. Thus, maladaptive UPR signaling through IRE1α may reinforce the transitional phenotype of these progenitor cells, exacerbating fibrosis while preventing normal lung repair.

## Results

### IRE1α acts on the pulmonary epithelium to promote fibrosis

We previously showed that a kinase inhibitor of IRE1α, KIRA7, protected mice from bleomycin when given starting on the day of bleomycin exposure through harvest.(22) We confirmed these results using a different kinase inhibitor based on an independent pharmacophore that we call KIRA8, which has nanomolar potency and high specificity for the IRE1α kinase.(23, 24) Bleomycin increased the size of lung fibrotic regions based on staining of fibrillar collagens I and III, and KIRA8 protected mice from these changes (Fig S1A).

The UPR has been shown to be activated in epithelial cells in human patients with IPF and mouse models of pulmonary fibrosis, suggesting that kinase inhibitors of IRE1α may protect from fibrosis in part by affecting epithelial stress signaling. To determine whether IRE1α activity is required in the epithelium for pulmonary fibrosis, we conditionally knocked out IRE1α using *Shh*^Cre^, which induces recombination throughout the lung epithelium from early development onward (Fig 1A). Loss of IRE1α activity in the epithelium was confirmed by reverse-transcription PCR for XBP1 mRNA splice isoforms in magnetically-enriched CD45^-^ EPCAM^+^ lung epithelial cells (Fig 1A). Mice were exposed to bleomycin and harvested on day 21. Conditional knockout of IRE1α in the epithelium protected mice from lung fibrosis based on wet weight (Fig 1B) and lung hydroxyproline content (Fig 1C), a highly specific measure of collagen, indicating that IRE1α activity in the epithelium is required for lung fibrosis.

**Figure 1.**
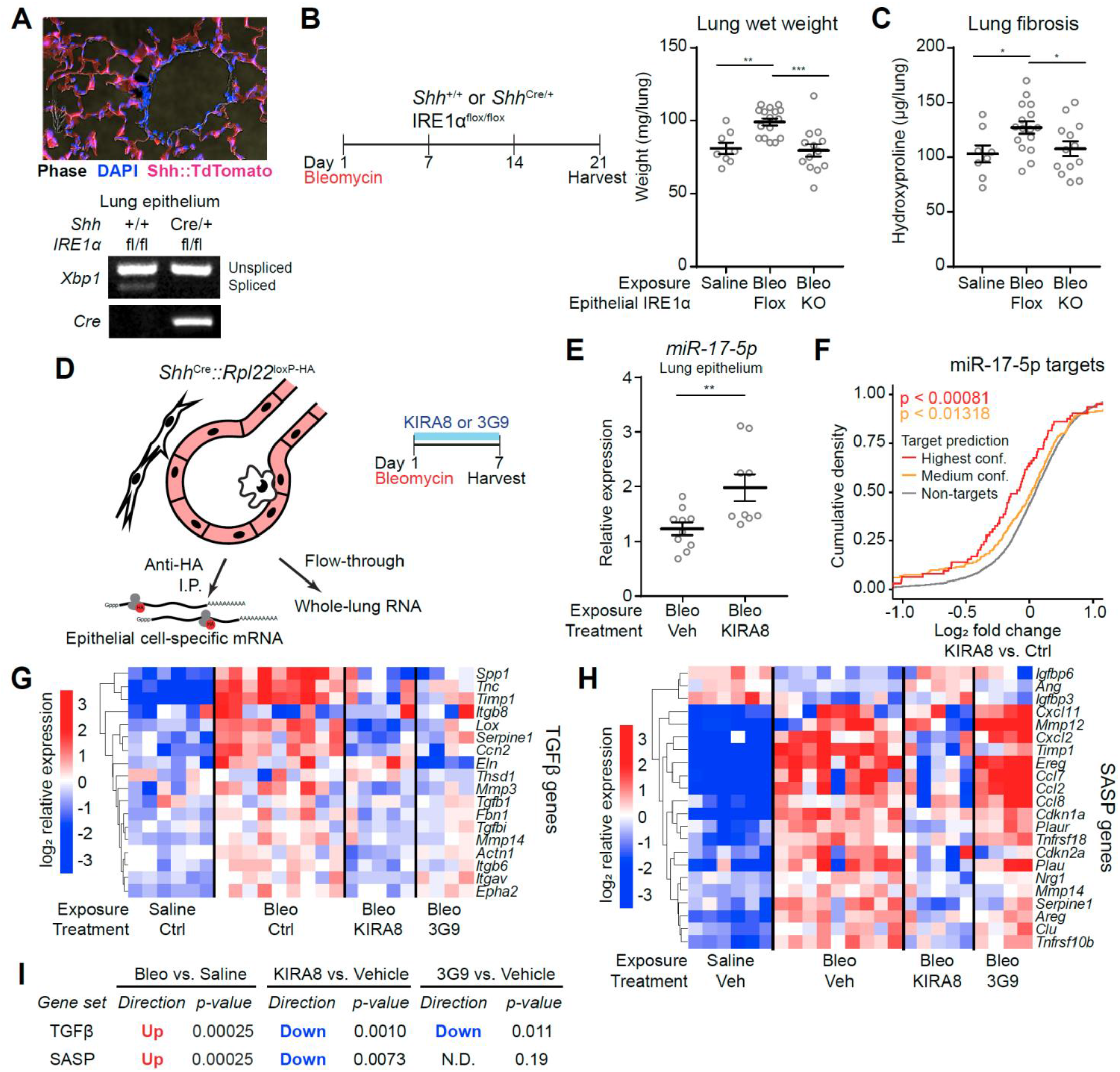
IRE1α regulates fibrosis and profibrotic gene expression in the lung epithelium. (A) Confirmation of Shh^Cre^ epithelial expression using *ROSA26*^Ai14^ reporter mice (top) and IRE1α loss of function (bottom). Lung epithelial cells were isolated by dissociation and MACS enrichment for CD45^-^ EPCAM^+^ cells, and IRE1α function evaluated by RT-PCR for XBP1 splicing. Presence of the Cre allele was confirmed by PCR. Mice were exposed to bleomycin, harvested at day 21, and left lung weight (B) and hydroxyproline (C) measured. Each data point represents one mouse, with mean ± SEM indicated. Groups were compared by ANOVA with Fisher’s post-hoc test. (D) RiboTag conditional ribosome tagging strategy. Lungs were harvested 7 days after bleomycin exposure and flash frozen. Epithelial ribosome-associated mRNA was purified by anti-HA immunoprecipitation. (E) Mature *mir-17-5p* expression in the lung epithelium after bleomycin exposure. Epithelial cells were isolated as in (A) and mature miRNA quantified by isoform-specific qPCR. Groups were compared using two-sided Student’s t-test. (F) Cumulative density function plot of batch- and sex-adjusted log_2_ fold changes for *miR-17-5p* target genes after KIRA8 treatment. *miR-17-5p* targets were obtained from TargetScan 7.2 and defined as high-confidence targets (percentile score > 99) or medium-confidence (percentile score between 95 and 99). Non-targets lack any *miR-17-5p* target site. Heatmaps of Batch- and sex-adjusted log_2_ expression values for TGFβ-induced (G) and SASP (H) genes in epithelial RiboTag libraries. SASP genes were filtered for those differentially expressed with FDR<0.1 between bleomycin- and saline-exposed epithelium. (I) Unfiltered gene set expression changes and statistical testing using the ROAST function in the limma package. * p<0.05, ** p<0.01, *** p<0.001.

To begin to understand how IRE1α promotes fibrosis in injured epithelial cells, we profiled the epithelial transcriptome on day 7 after bleomycin exposure, a timepoint when markers of ER stress are first elevated in whole lung RNA(22) and when gene expression programs in injured epithelial cells, like TGFβ signaling, begin to establish the fibrotic milieu. We used RiboTag mice, in which the ubiquitous ribosomal protein Rpl22 is conditionally labeled with the HA epitope by Cre recombinase.(25) This tool enables immunoprecipitation of cell-type specific mRNAs from flash-frozen lungs and reflects both transcriptional abundance and ongoing translation, while avoiding gene expression artifacts that have been described in dissociated and sorted cells, especially in stress signaling pathways.(26) *Shh*^Cre^ was used to label all epithelial subtypes in the lung (Fig 1D, Fig S1B) after bleomycin exposure and treatment with either KIRA8, an inhibitor of the IRE1α kinase, or 3G9, an antibody that blocks integrin αvβ6 and TGFβ signaling.(27)

Hyperactivation of IRE1α can cause epithelial cell dysfunction through several proposed mechanisms. One is IRE1α phosphorylation of c-Jun N-terminal kinase (Jnk).(28) Consistent with on-target inhibition of the IRE1α kinase, KIRA8 downregulated genes in the Jnk pathway in the lung epithelium by Ingenuity Pathway Analysis (Fig S1C). IRE1α has also been shown to degrade ER-localized RNAs, a process called Regulated IRE1α-dependent decay (RIDD). *Bloc1s1* (Blos1) is a well-characterized RIDD substrate of IRE1α in multiple cell types.(3, 4, 29) Inhibiting IRE1α with KIRA8 alleviated downregulation of *Bloc1s1* in the lung epithelium, but did not affect *Bloc1s1* expression in the whole-lung transcriptome (Fig S1D). RIDD activity has recently been shown to enhance activation of DNA damage pathways.(16) Consistent with this observation, KIRA8 treatment dampened the *Tp53* (encoding p53) and *Ddx5* (encoding p68) pathways in the lung epithelium after bleomycin exposure (Fig S1C).

In addition to ER-localized mRNAs, hyperactivated IRE1α RIDD degrades select microRNAs (miRNAs) precursors such as pre-miR-17, thus leading to decreases in mature miR-17, which has been shown to de-repress *Txnip* and promote apoptosis.(5, 30) Importantly, decreases in miR-17 have been described in lungs from both IPF patients and mice exposed to bleomycin, and miR-17 has been implicated in stem cell regulation, senescence, and TGFβ signaling, all important to IPF.(31–36) We reasoned that inhibiting IRE1α would increase mature miR-17-5p and consequently increase repression of its mRNA targets. Indeed, in magnetically-enriched CD45^-^ EPCAM^+^ lung epithelial cells, KIRA8 treatment increased the abundance of mature miR-17-5p. In the epithelial RiboTag data, KIRA8 correspondingly increased repression of conserved miR-17-5p targets defined in TargetScan 7.2, a validated tool(37) that accurately predicts targets of miRNA repression (Fig 1F).

To further understand how epithelial IRE1α regulates fibrosis, we evaluated epithelial pathways known to play a role in fibrosis. One pathway is TGFβ, a key activator of collagen-producing fibroblasts. *In vitro*, UPR activation has been shown to trigger TGFβ signaling and vice versa, including through IRE1α’s RIDD activity.(20, 38–40) *In vivo*, ER stress, selectively induced in the lung epithelium, causes increases in TGFβ signaling.(8, 9) We used microarray data from bleomycin-exposed *Itgb6* knockout mice to define a signature of TGFβ-induced genes.(41) Integrin αvβ6 is expressed on injured lung epithelial cells and releases active TGFβ from the extracellular matrix, which in turn causes autocrine induction of *Itgb6* transcription and other TGFβ-induced genes. In our epithelial RiboTag data, these genes were upregulated after bleomycin exposure. Remarkably, KIRA8 downregulated these genes to a similar degree compared to 3G9, a direct inhibitor of αvβ6-mediated TGFβ activation (Fig 1G and I). These results suggest that the UPR, and specifically activated IRE1α, are upstream of activation of the αvβ6—TGFβ axis.

Pulmonary fibrosis is not a uniform process, but rather develops in discrete, localized regions. These regions are characterized by close association of dysmorphic lung epithelial cells and activated fibroblasts that produce excessive amounts of collagens and other extracellular matrix proteins, perhaps in response to epithelial signals. Multiple recent reports have described dysfunctional epithelial cells that could contribute to this pathologic conversation. These cells are characterized by a phenotype reminiscent of cellular senescence and upregulation of a gene expression signature called the “senescence-associated secretory phenotype” (SASP), although the signature is not specific to senescence and may be a general indicator of epithelial dysfunction.(42–44) This signature is upregulated in epithelial cells after bleomycin injury and decreased after KIRA8 treatment (Fig 1H and I). One key marker in this signature is *Cdkn1a*, which encodes p21, a cyclin dependent kinase inhibitor that induces cell cycle arrest and is a canonical marker of senescent cells. *Cdkn1a* was induced by bleomycin exposure and this was blunted by KIRA8 treatment. Although TGFβ signaling can by itself induce senescence, 3G9 did not mitigate either *Cdkn1a* upregulation or the SASP signature as a whole (Fig 1H and I), indicating that IRE1α independently regulates TGFβ signaling and the SASP, and that TGFβ signaling is not required for epithelial SASP gene expression.

### Damage-associated transient progenitor cells (DATPs) express IRE1α-regulated profibrotic gene expression programs

A parsimonious explanation of these findings would be that IRE1α regulates a single subpopulation of lung epithelial cells after bleomycin exposure, inducing loss of miR-17-5p and consequent upregulation of miR-17 targets, TGFβ signaling, and SASP gene expression. We re-analyzed a recently published single-cell RNA sequencing dataset from bleomycin exposed mice in which lung epithelial cells were sampled at daily intervals after bleomycin exposure (Fig 2A).(45) All three gene expression signatures blocked by KIRA8—TGFβ signaling, SASP gene expression, and upregulation of miR-17 targets—were highest in a discrete population of progenitor cells in a “transitional” state that expand dramatically after bleomycin exposure (Fig 2B), called damage-associated transient progenitor (DATP) cells. Several recent studies have demonstrated that these cells emerge after multiple types of lung injury, including bleomycin exposure, epithelial cell ablation by diphtheria toxin, and viral infection; are derived from both airway and alveolar epithelial cells; co-express multiple cytokeratins (*Krt7, Krt8*, and *Krt19*, Fig 2C); and are capable of trans-differentiating into mature type 1 and type 2 alveolar epithelial cells (AEC1 and AEC2).(45–49) After bleomycin injury, these cells exhibit a dysfunctional phenotype with reduced capacity to proliferate and differentiate, and instead express genes that have been associated with fibrosis.

**Figure 2.**
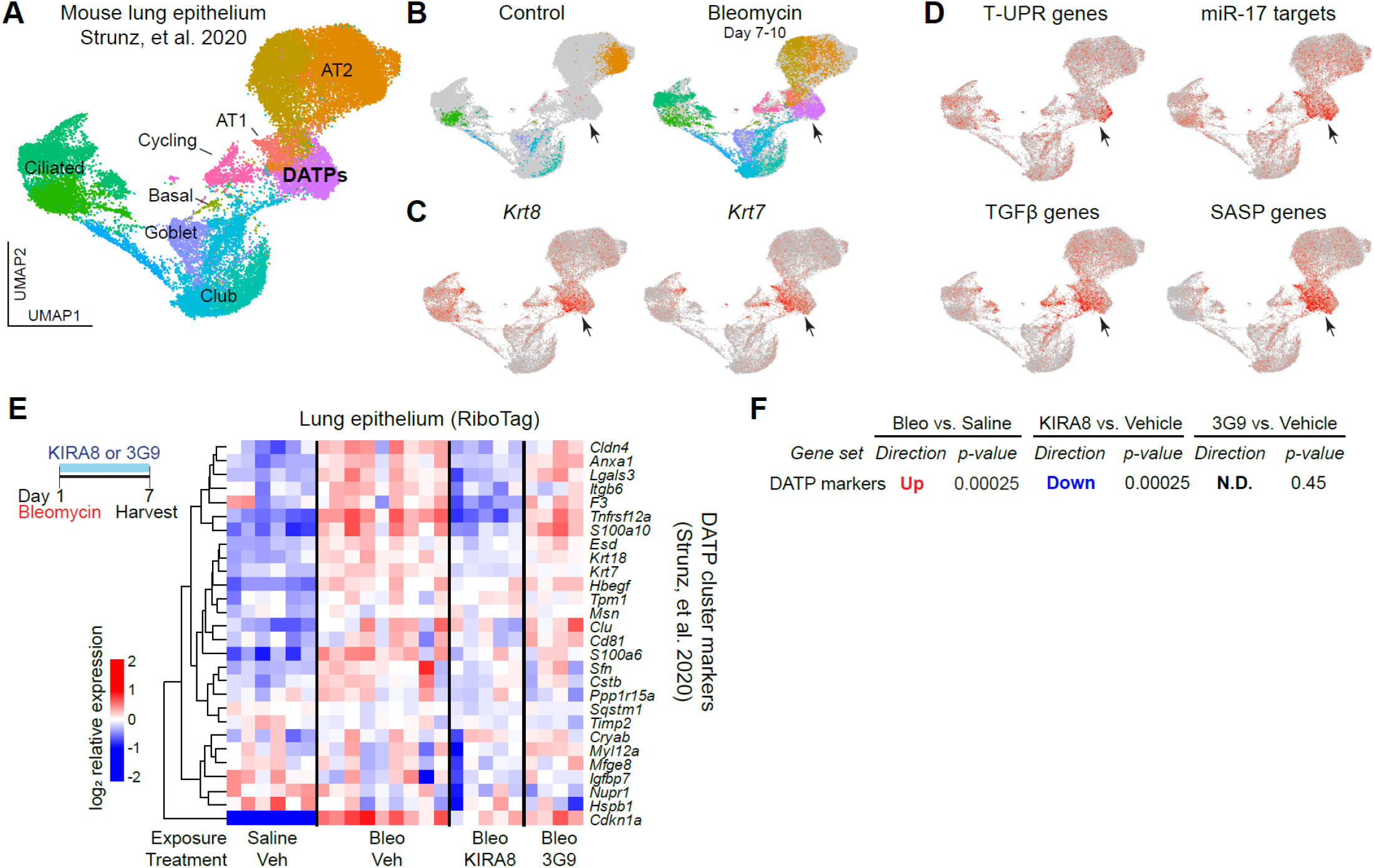
IRE1α-regulated gene expression signatures are preferentially expressed in damage-associated transient progenitor (DATP) cells. (A) Single cell RNA sequencing of bleomycin-exposed lung epithelium. The UMAP projection and cluster identities were defined previously.^45^ (B) UMAP projections of subsets of the dataset in (A) for cells from control lungs (left) and lungs harvested day 7 through day 10 after bleomycin exposure (right). Gray denotes cells not in the respective subset; cluster colors and identities are as in (A). (C) Overlay of *Krt8* and *Krt7* expression. (D) Overlay of AUCell scores for gene signatures regulated by IRE1α. Arrowheads denote the DATP population. (E) Heatmap of markers of the DATP cluster in the lung epithelium after bleomycin exposure and treatment with KIRA8 or 3G9. Markers of were defined by the Seurat package using single cell RNA sequencing data in ^45^. (F) Unfiltered DATP marker gene expression changes and statistical testing using the ROAST function in the limma package.

Consistent with recent descriptions of dysfunction, we found that DATPs upregulated genes associated with pathological or ‘terminal’ UPR (T-UPR) signaling during their emergence (Fig 2D). To evaluate whether UPR signaling through IRE1α indeed regulated DATPs, we examined our epithelial RiboTag data for the expression of genes uniquely expressed in this cluster of cells. Inhibition of the IRE1α kinase with KIRA8 decreased expression of genes marking these cells (Fig 2E and F). Although one study recently proposed that TGFβ might regulate these cells,(48) we did not find the same effect with the anti-integrin αvβ6 antibody 3G9 (Fig 2E and F), suggesting that if TGFβ does regulate induction of these cells in vivo, TGFβ is activated by an αvβ6-independent mechanism.

Our findings suggested that IRE1α might regulate the number or the dysfunctional phenotype of DATPs, including activation of TGFβ regulated genes. To identify DATPs in lung sections, we used antibody staining for Krt8 protein or in situ hybridization for *Krt7* mRNA. At the transcriptional level, *Krt7* mRNA is a more faithful marker for DATPs because *Krt8* is expressed at low levels in normal ciliated bronchial epithelial cells (Fig 2C and Fig 3A). Consistent with previous studies, *Krt7+* DATP number increased dramatically after bleomycin exposure. This increase was blocked by KIRA8 (Fig 3B). We evaluated apoptosis of these cells at day 7, when they are near their peak numbers.(45) KIRA8 treatment did not cause widespread apoptosis of these cells, since TUNEL+ Krt8+ double-positive cells were exceptionally rare in both vehicle and KIRA8 treated mice (Fig S2). Thus, IRE1α signaling either promotes induction or maintenance of this population.

**Figure 3.**
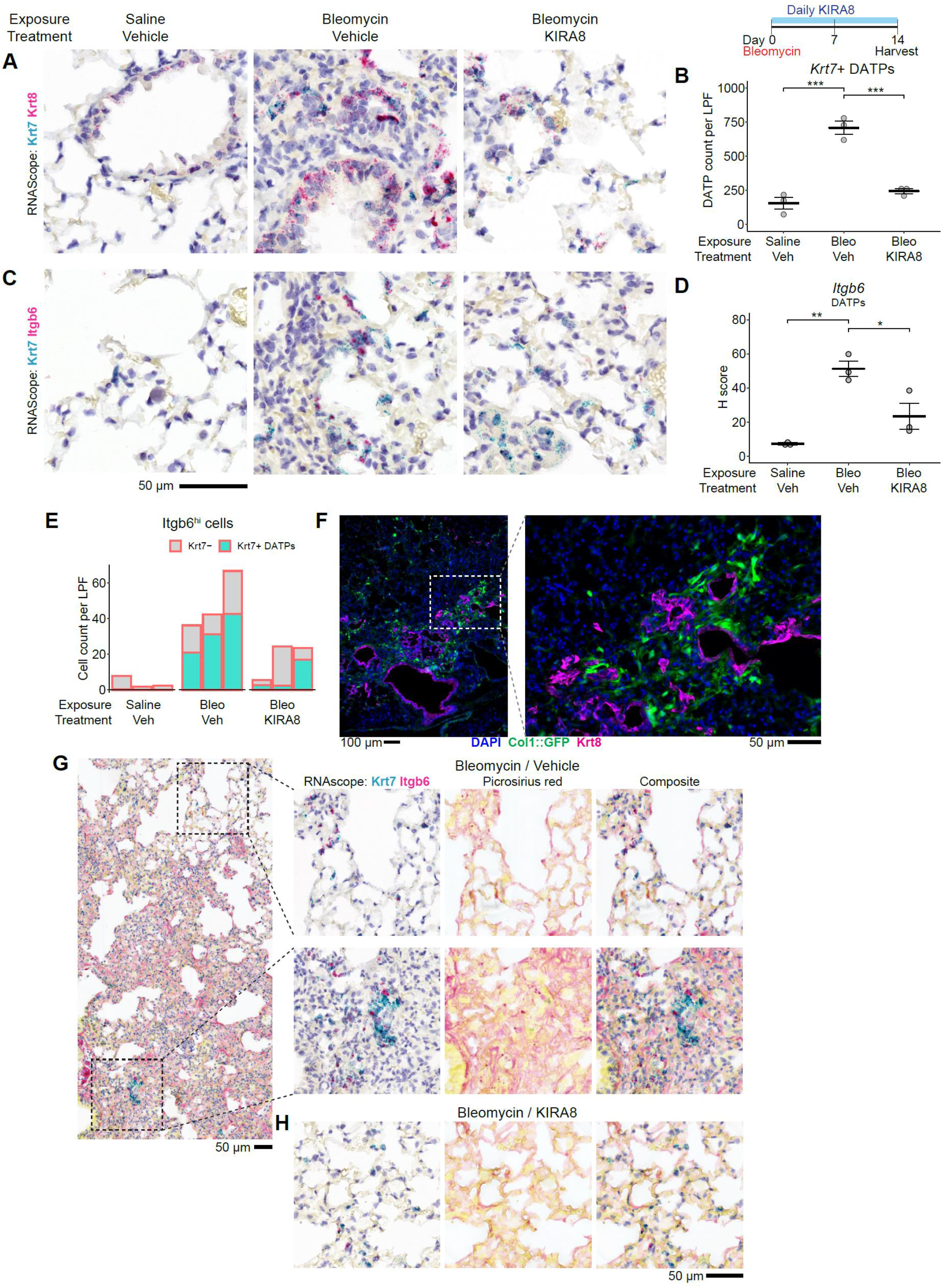
DATPs are situated in the fibrotic niche and are regulated by IRE1α. (A) RNA in situ hybridization (RNAscope) for *Krt7* (teal) and *Krt8* (magenta) at day 14 after bleomycin exposure and treatment with KIRA8. (B) Automated count of *Krt7*+ cells per 1240 µm diameter low-power field (LPF). Each dot represents the average of 10 representative fields from one mouse. Groups were compared by ANOVA with Fisher’s post-hoc test. (C) RNA in situ hybridization for *Krt7* (teal) and *Itgb6* (magenta) as in (A). (D) Semi-quantitative H-score of *Itgb6* puncta in *Krt7*+ DATPs, averaged over 10 representative fields per dot as in (B). (E) Distribution of *Itgb6*^high^ cells among *Krt7*+ or *Krt7*-cells, averaged over 10 representative fields per bar as in (B). (F) Representative field of Krt8 immunofluorescence staining (magenta) in a section from bleomycin-exposed *Col1*-GFP mice at day 14. (G) RNA in situ hybridization for *Krt7* (teal) and *Itgb6* (magenta) as in (A), imaged, cleared and re-stained for fibrillar collagen with picrosirius red.

### IRE1α regulates TGFβ activation by DATPs

Though KIRA8 decreased DATP number, we nonetheless found clusters of *Krt7*+ DATPs in regions of injured lungs after KIRA8 treatment and wondered whether IRE1α regulates the profibrotic phenotype of the remaining cells. In pulmonary fibrosis, epithelial expression of integrin αvβ6 activates TGFβ from the extracellular matrix and signals to nearby fibroblasts to mature and produce collagen. Since KIRA8 substantially dampened TGFβ signaling (Fig 1G and I), we evaluated the integrin αvβ6—TGFβ axis in the progenitor population by RNA in situ hybridization (Fig 3C). We found that the majority of *Itgb6* expressing cells in the bleomycin-exposed lungs were marked by *Krt7* mRNA (Fig 3C and 3E). KIRA8 treatment decreased the expression of *Itgb6* in *Krt7+* DATPs (Fig 3C and 3D).

We reasoned that since dysfunctional progenitor cells appeared to be the principal source of integrin αvβ6-induced TGFβ activation, they would be co-localized with activated fibroblasts marked by high levels of collagen expression. Though *Krt7* is a more faithful mRNA marker, we used Krt8 as a protein marker due to the availability of well-characterized antibodies. We found that after bleomycin exposure Krt8+ DATPs were surrounded by fibroblasts with high collagen 1 expression; conversely, activated fibroblasts were sparse in regions without Krt8+ cells (Fig 3F). Correspondingly, DATPs marked by *Krt7* RNA were surrounded by extracellular collagen stained with picrosirius red (Fig 3G). Consistent with its effect on *Itgb6* expression and downstream TGFβ signaling, KIRA8 treatment decreased the density of picrosirius red around Krt8+ progenitors (Fig 3H).

### IRE1α regulates dysfunctional features of DATPs

IRE1α may also regulate fibrosis through its effects on “SASP” gene expression, since chemical or genetic ablation of cells with this phenotype has been shown to protect against fibrosis.(44, 50) IRE1α upregulation of these dysfunctional epithelial gene programs may impact the ability of DATPs to proliferate and differentiate into mature epithelial cells, even if these cells are not senescent in the sense of permanent cell cycle arrest. In our RiboTag sequencing, *Cdkn1a* expression was increased in epithelial cells after bleomycin exposure, and this increase was blunted by KIRA8 inhibition of IRE1α (Fig 1H and 4A). Likewise, we found *Cdkn1a* expression increased in DATPs after bleomycin exposure (Fig 4B and 4C), consistent with previous reports finding reduced proliferative capacity in these cells after injury.(45) KIRA8 decreased *Cdkn1a* expression in *Krt7+* DATPs (Fig 4B and C), suggesting that IRE1α signaling might exacerbate the dysfunctional phenotype of these cells. In addition, after KIRA8 treatment, we found rare clusters of Krt7+ cells integrated in alveolar structures that expressed *Ager* (RAGE), a marker of mature AEC1s (Fig S3). These dual-positive clusters were previously suggested to be progenitor cells in the process of differentiating into mature epithelium.(45) Though dual *Krt7+ Ager*+ cells are present in both vehicle and KIRA8 treated mice, we did not find any such cells integrated in alveoli in vehicle-treated mice. Together, these results suggest that that inhibiting IRE1α may decrease the apparent progenitor cell count in part by promoting their proliferation and differentiation into mature AECs.

**Figure 4.**
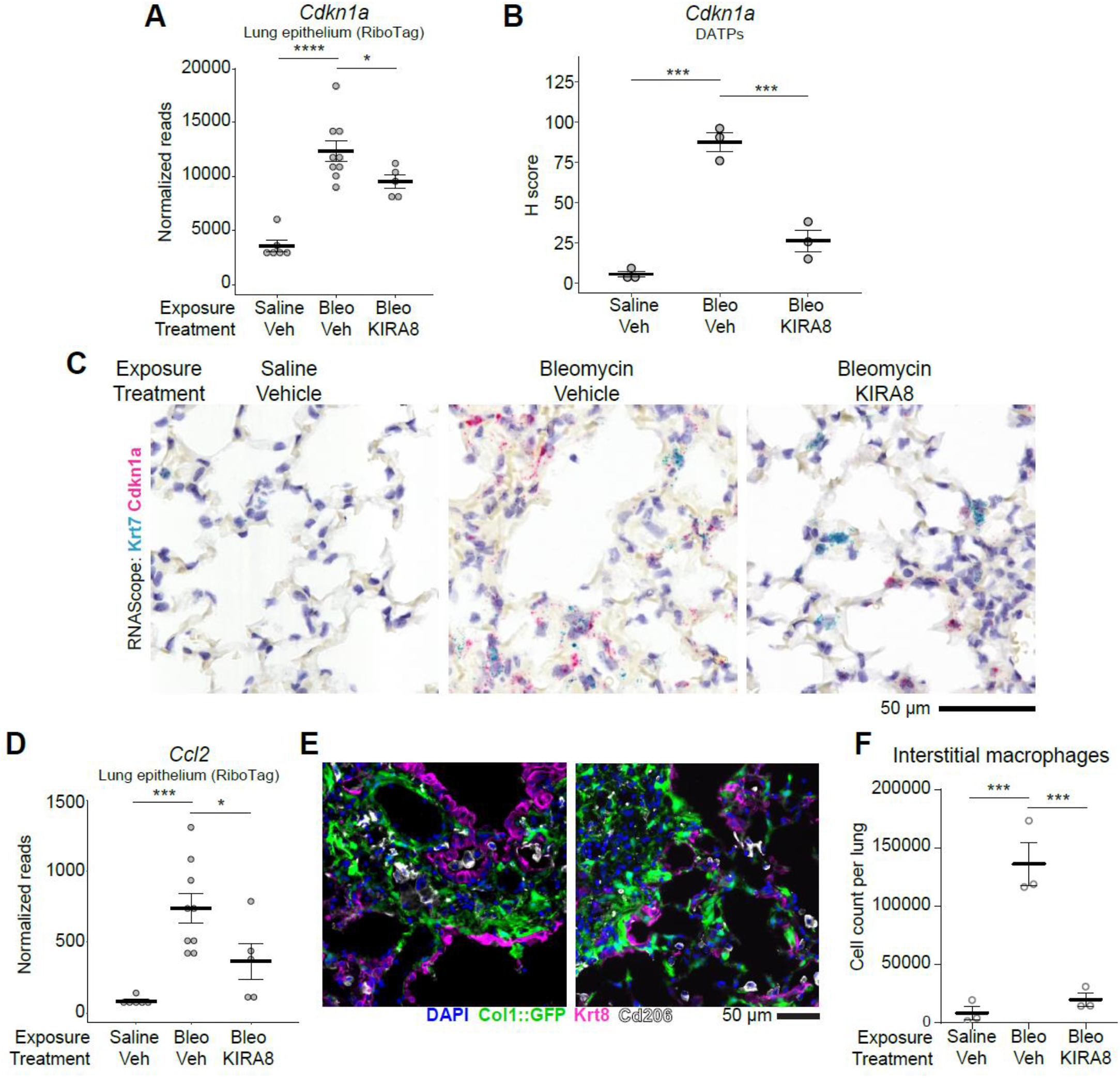
Senescence-associated phenotypes of DATPs are regulated by IRE1α. (A) *Cdkn1a* expression in epithelial RiboTag data on day7 after bleomycin exposure and treatment with KIRA8. Read counts were adjusted for batch and sex. Groups were compared by ANOVA with Fisher’s post-hoc test. (B) Semi-quantitative H-score of *Cdkn1a* puncta in DATPs, as in Figure 3D. (C) RNA in situ hybridization (RNAscope) for *Krt7* (teal) and *Cdkn1a* (red) as in Figure 3A. (D) *Ccl2* expression in epithelial RiboTag data, as in Fig 4A. (E) Representative field of Krt8+ (magenta) and CD206 (white) immunofluorescence staining in a section from bleomycin-exposed mice at day 14. (F) Flow cytometric quantitation of interstitial macrophages after bleomycin exposure and treatment. Lungs were dissociated to single cell suspension and macrophages were defined as in ^72^ as CD45^+^ CD24^-^ F4/80^+^ CD11b^high^ Ly6C^low^ CD11c^+^ MHC-II^high^. Groups were compared by ANOVA with Fisher’s post-hoc test. * p<0.05, ** p<0.01, *** p<0.001, **** p<0.0001.

Several genes in the “SASP” signature expressed by DATPs after bleomycin exposure (and downregulated by KIRA8) are well-known monocyte chemoattractants and monocyte activators, including *Ccl2* (MCP-1), *Ccl8* (MCP-2), *Ccl7* (MCP-3), *Ccl3* (MIP-1α), *Cxcl2* (MIP-2α) (Fig 1H and 4D). Recent studies have described monocytes infiltrating the lung after bleomycin injury, becoming CD11b^high^ CD11c^+^ MHC-II^high^ “interstitial” macrophages over several days, and ultimately differentiating over weeks into monocyte-derived alveolar macrophages. These newly-recruited macrophages have been proposed to help establish the fibrotic niche by expressing fibroblast growth factors such as PDGFα.(51–54) In turn, IL-1β released by these macrophages can prevent DATP differentiation, instead trapping them in the dysfunctional state.(49) On day 14 after bleomycin exposure Krt8+ DATPs were co-localized with macrophages marked by CD206, a marker that is broadly expressed on macrophages (Fig 4E). Lung macrophages at this time point after bleomycin have been shown to be predominantly monocyte-derived,(55) suggesting that DATPs can help recruit these players to the fibrotic niche. Since inhibiting IRE1α with KIRA8 downregulated lung epithelial expression of monocyte chemoattractants on day 7 after bleomycin exposure (Fig 4D), we reasoned that KIRA8 treatment would cause decreases in infiltrating interstitial macrophages. Indeed, CD11b^high^ CD11c^+^ MHC-II^high^ interstitial macrophages increased in bleomycin exposed lungs, and this increase was abolished by KIRA8 treatment (Fig 4F).

### DATP analogs in IPF lungs express IRE1α-regulated profibrotic transcriptional programs

To determine whether a similar phenomenon occurs in human IPF lungs, we combined single-cell RNA sequencing of epithelial cells from 1 healthy donor lung and 4 IPF lungs with a public dataset including an additional 10 IPF lungs and 6 normal lungs (GSE135893; (56)) (Fig 5A). Two discrete epithelial populations were found exclusively in diseased lungs and expressed all the gene expression signatures associated with the T-UPR, TGFβ signaling, SASP gene expression, and upregulation of miR-17 targets (Fig 5B and 5C). Both these populations are marked by KRT17, including a recently described group of KRT17+, KRT5-cells (Fig 5B)(57) that are proposed to be the human equivalents of mouse DATPs.(45) Ligand-receptor analysis revealed multiple potential intercellular signals emanating from KRT17+ KRT5-cells to fibroblasts and monocyte-derived macrophages (Fig 5D). These results suggest that inhibiting IRE1α in human IPF may ameliorate the profibrotic phenotype of dysfunctional progenitor cells, as it does in the bleomycin model.

**Figure 5.**
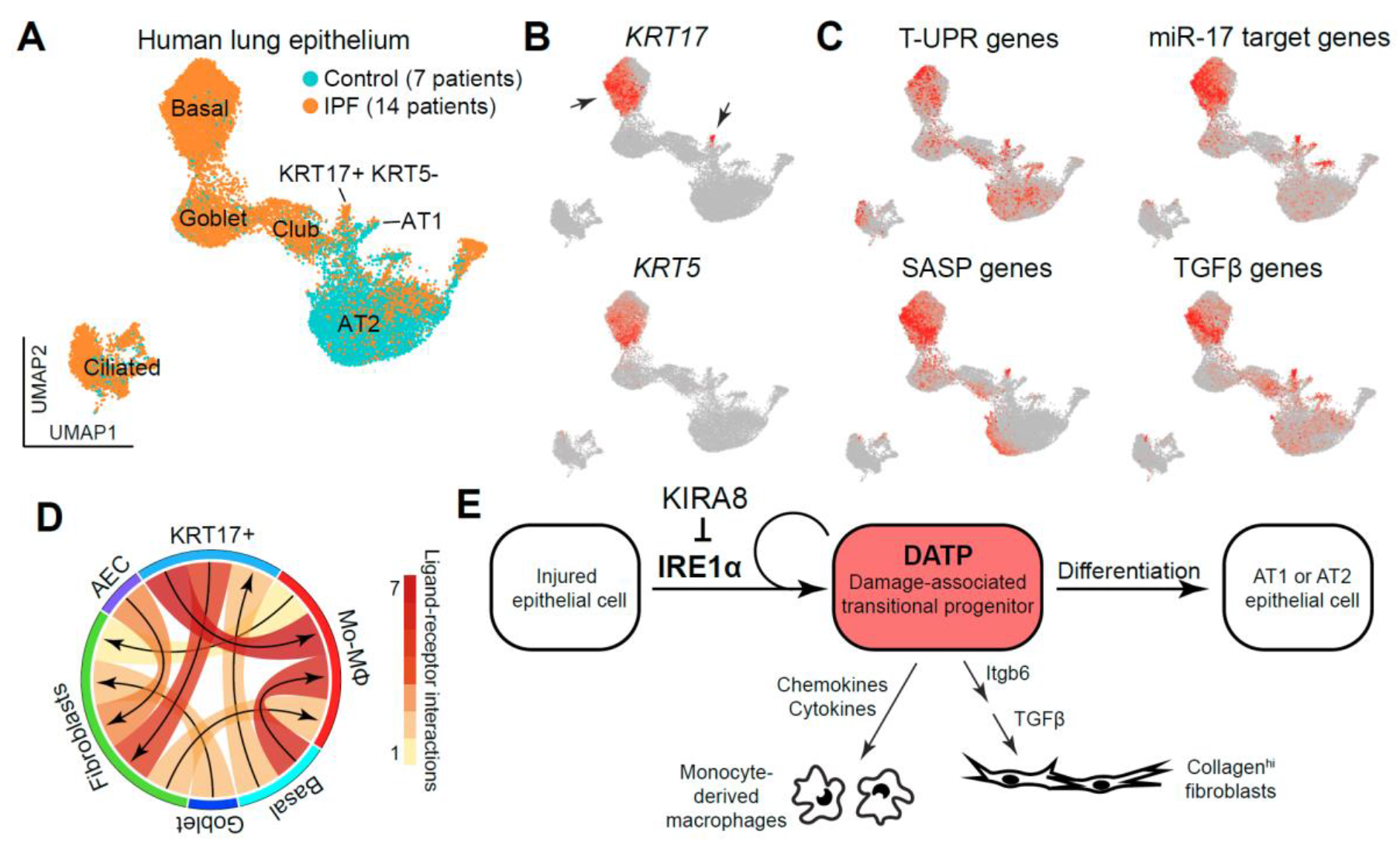
Basal cells and *KRT5*-*KRT17*+ “basaloid” epithelial cells in human IPF lungs have analogous gene expression signatures as mouse DATPs. (A) UMAP dimensional reduction plot of single cell sequencing of the epithelium from IPF patients or donor controls. (B) Overlay of *KRT5* and *KRT17* expression in the human lung epithelial dataset. (C) Overlay of AUCell scores in the human lung epithelial dataset for gene signatures that are regulated by IRE1α in the mouse lung. (D) Ligand-receptor expression analysis of interactions between lung epithelial cells, fibroblasts, and monocyte-derived macrophages in the human single-cell dataset. Arrows denote ligand-receptor pairs with arrows pointing from ligand towards receptor. (E) Model for IRE1α regulation of fibrosis in injured lung epithelial cells. IRE1α either mediates conversion of injured epithelial cells into DATPs, or helps maintain the dysfunctional phenotype. In the dysfunctional state, DATPs express profibrotic factors including *Itgb6* that releases TGFβ to activate fibroblasts, and “SASP” chemokines to recruit profibrotic monocyte-derived macrophages.

## Discussion

ER stress and consequent maladaptive activation of UPR pathways have been observed in epithelial cells in IPF and other fibrotic lung diseases.(6, 7, 11, 58) One such switch between adaptive and maladaptive outcomes in the UPR is mediated by the ER transmembrane kinase/RNase, IRE1α. We previously showed that interrupting the IRE1α arm of the UPR using highly potent and selective kinase inhibitors, which allosterically inhibit its RNase catalytic activity, protected mice from bleomycin-induced fibrosis.(22) In this study, we unexpectedly found that one such inhibitor, KIRA8, modulated the number and dysfunctional phenotype of damage-associated transient progenitor (DATP) cells. Using conditional ribosome tagging of the epithelial transcriptome, we found that KIRA8 treatment increased mature miR-17 levels, and dampened TGFβ signaling and senescence-associated gene expression. In single-cell RNA sequencing analysis, we found these programs were selectively expressed in DATPs. In fact, these cells were the predominant source of *Itgb6* expression, and were correspondingly localized to fibrotic regions, surrounded by pathological fibroblasts and macrophages. Inhibiting IRE1α signaling decreased *Itgb6* expression and surrounding fibrosis, and decreased the infiltration of profibrotic interstitial macrophages. We observed the same gene expression signatures in KRT17+ “basaloid” cells in the lungs of IPF patients, suggesting that these results may be applicable to human disease.

An opposing dynamic has long been recognized between lung regeneration and fibrosis. Several recent papers have described a novel population of progenitor cells in the lung that emerge after injury.(45–48) These cells mirror this dynamic as they progress from injured mature epithelium into de-differentiated “transitional” cells (DATPs) and then differentiate again into mature epithelium to repair the lung. While in the transitional state, DATPs enact gene expression programs regarded as pathological or dysfunctional, including those associated with hypoxia, DNA damage, senescence, ER stress, and fibrosis.(45–48) Intriguingly, some of these programs, such as the p53 pathway, may actually be required for subsequent differentiation into mature epithelium.(47) This suggests that, in DATPs, the tension between regeneration and fibrosis resides in the decision to abnormally remain in the transitional state, rather than proceed with normal differentiation and repair.(59, 60)

Our results suggest that cellular stress signaling through IRE1α may contribute to the stalling of DATPs in the dysfunctional/transitional state, along with other cell-intrinsic programs and extrinsic cues. Trapped in this state, DATPs are central players in establishing the fibrotic niche and concomitantly unable to execute normal lung repair programs. Inhibiting IRE1α signaling decreases profibrotic gene expression in the cells, including expression of integrin β6, and decreases expression of chemokines that recruit profibrotic monocyte-derived macrophages. Inhibiting IRE1α signaling may also restore proliferation and differentiation capacity, a question that will be interesting to address with newly-described lineage tracing tools.(47)

ER stress and activation of the UPR in the lung epithelium, caused by loss of protein folding capacity or a high burden of misfolded ER luminal protein, can be sufficient to cause pulmonary fibrosis in both mouse models and in human disease.(7–9) The UPR is also secondarily activated in DATPs across disparate models of lung injury,(45) but the precise cause of this activation is not clear and may be multifactorial. DNA damage signaling, senescence, hypoxia and inflammation have all been shown to particularly activate IRE1α, even in the apparent absence of unfolded protein, and these triggers are deeply conserved in evolution.(12, 13, 15, 16, 61–63) Our results are reminiscent of findings in induced pluripotent stem cells (iPSCs), where transient activation of the IRE1α pathway enhances reprogramming of fibroblasts into iPSCs, but sustained activation over days causes aberrant morphology, hypoproliferation, and eventually stem cell loss.(64)

The root of IRE1α activation is part of a broader mystery of how or why a broad-spectrum cellular stress response is engaged in these cells. For example, the p53 pathway is activated in DATPs even when the primary lung injury would not be expected to cause DNA damage,(47, 48) and despite its necessity, the mechanistic role of p53 in DATP differentiation is unclear. One downstream role for IRE1α activation may be its degradation of the miR-17 precursor, leading to upregulation of miR-17 target genes.(5) Indeed, we observed upregulation of mature miR-17-5p after KIRA8 inhibition of IRE1α, with corresponding downregulation of its target mRNAs (Fig 1), which include genes important to TGFβ signaling and senescence, such as *Tgfbr2*, encoding a subunit of the TGFβ receptor;(65) *Ctgf*, a TGFβ-induced fibroblast activator;(66) and *Cdkn1a*, encoding p21.(67) Indeed, miR-17 overexpression enables proliferation of lung epithelial progenitors in early development,(31) its loss is embryonic lethal due to lung hypoplasia,(68) and it is strikingly downregulated in IPF.(69) A deeper understanding of cellular stress response pathways in progenitor cells may ultimately enable therapies—such as kinase inhibition of IRE1α—that both ameliorate fibrosis and enhance regeneration in human lung disease.

## Methods

### Mice

All mice used were C57BL/6 mice or congenic to the C57BL/6 background. Wildtype mice were obtained from Jax (000664). Transgenic mice were: *Shh*^CreGFP^ mice (Jax 005622), RiboTag mice (Jax 011029), *Ern1*^flox/flox^ (gift of T. Iwawaki), Tg(Col1a1-EGFP) (gift of D.A. Brenner). All work was approved by the Institutional Animal Care and Use Committee of the University of California, San Francisco. To induce fibrosis, 8-14 week old mice were anesthetized with either isoflurane or ketamine and xylazine and exposed to a single dose of intranasal bleomycin (3 units/kg). Mice were sacrificed by open-drop isoflurane overdose and lungs were harvested at the indicated times. KIRA8 was dissolved in a vehicle consisting of 3% ethanol, 7% Tween-80, and 90% normal saline and injected peritoneally at 50 mg/kg daily starting on the day of bleomycin exposure. The anti-integrin β6 antibody 3G9 or the inert control antibody AXUM8 was dissolved in normal saline and injected peritoneally at 10 mg/kg twice a week starting on the day of bleomycin exposure.

For total hydroxyproline content, lungs were processed in a rotor-stator homogenizer, protein precipitated with tricholoracetic acid, and the pellet boiled in concentrated hydrochloric acid overnight. Hydroxyproline was oxidized using chloramine T and reacted with Ehrlich’s solution and color change measured by absorbance at 550 nm.

### Histology

Formalin-fixed, paraffin embedded sections were prepared by inflating lungs with zinc formalin fixative, incubating in formalin overnight followed by 70% ethanol, then paraffin embedding and sectioning. Frozen sections were prepared by inflating lungs with 1.5% paraformaldehyde in OCT, incubating in 4% paraformaldehyde for 1 hour, washing in PBS for 1 hour, serial flotation in 30% sucrose/PBS and 15% sucrose/OCT, embedding in OCT and sectioning. Picrosirius red (Abcam ab150681) or TUNEL (Invitrogen C10617) staining was performed per the respective manufacturers’ instructions. RNAscope staining (ACD Bio) was performed on formalin-fixed, paraffin-embedded sections by the manufacturer using the following probes: Mm-Krt7-C1 (511808), Mm-Krt8-C2 (424521), Mm-Itgb6-C2 (312501), Mm-Cdkn1a-C2 (408551), and Mm-Ager-O1-C2 (550798). Antibody staining was performed using the following primary antibodies: rabbit anti-Krt8 (Abcam EPR1628Y, 1:500), rat anti-CD206 (BioRad MCA2235GA, 1:200). For quantification, ten representative low-power fields (1240 µm in diameter) were extracted from one section for each mouse lung, and quantified for picrosirius red staining area or RNA puncta counts and associated with cells defined by hematoxylin stained nuclei using the CellProfiler package.

### Epithelial cell isolation and reverse-transcription PCR

Epithelial single cell suspensions were prepared as described previously.(70) Magnetic nanoparticle depletion and enrichment was performed MACS (Miltenyi). Cells were depleted for CD16/32 (Miltenyi 130-101-895) and CD45 (130-052-301), followed by positive selection for EPCAM (Miltenyi 130-101-859). Total RNA was prepared using Tri-Reagent (Ambion). For XBP1 splice isoform detection, cDNA was synthesized using the Quantitect Reverse Transcription Kit (Qiagen), followed by PCR using the primers ACACGCTTGGGAATGGACAC and CCATGGGAAGATGTTCTGGG and separation on 4% agarose. For mature miRNA quantification, target-specific reverse transcription was performed using the TaqMan MicroRNA Reverse Transcription Kit (Thermo) followed by quantitative PCR with the TaqMan Universal Master Mix (Thermo), using primer sets for *mir-17-5p* (Thermo 002308) and *snoRNA202* (Thermo 001232).

### RiboTag immunoprecipitation

Lungs from *Shh*^Cre^ *Rpl22*^flox/flox^ mice were dissected and flash-frozen in liquid nitrogen within a few minutes of respiratory arrest and stored at −80° C. All immunoprecipitation steps were performed at 4° C. Lungs were thawed into 10 volumes of Supplemented Homogenization Buffer (100 mM KCl, 50 mM Tris-Cl pH 7.4, 12 mM MgCl_2_, 1% NP-40, 1 mM DTT, 0.1 mg/ml cycloheximide, 1.25 mg/ml heparin, 400 U/ml NEB murine RNase inhibitor) and dounced. The lysate was cleared by centrifugation and an aliquot taken for whole lung RNA preparation using the RNeasy Mini kit (Qiagen). The remaining lysate was incubated for 4 hours with a 1:20 dilution of anti-HA magnetic beads (Pierce 88837). Beads were washed once in Supplemented Homogenization Buffer, transferred to a fresh tube, then washed three times in High Salt Wash Buffer (300 mM KCl, 50 mM Tris-Cl pH 7.4, 12 mM MgCl_2_, 1% NP-40, 0.5 mM DTT, 0.1 mg/ml cycloheximide, 80 U/ml NEB murine RNase inhibitor). Beads were transferred to a fresh tube and immunoprecipitated RNA extracted by incubating beads in buffer RLT (Qiagen) supplemented with 2-mercaptoethanol and purified using the RNeasy Micro kit (Qiagen).

### RNA sequencing and data analysis

Libraries were prepared for sequencing by the UCSF Genomics CoLab using the Quantseq Forward kit (Lexogen) and sequenced on a HiSeq 4000 (Illumina). Reads were aligned to the mouse genome (Ensembl GRCm38) using STAR 2.5.2b. Differential expression testing was performed by the DESeq2 package using the model ∼batch + sex + treatment group. For heatmaps, expression data was transformed using the rlog function in DESeq2 and batch-corrected using the removeBatchEffect function in the limma package. Fold-change results from DESeq2 were exported to Ingenuity Pathway Analysis (Qiagen) for upstream regulator analysis. Custom gene set statistical testing was performed using the ROAST function in the limma package. miRNA targets and percentile scores were extracted from TargetScan 7.2; non-targets lacked any predicted site, regardless of conservation. Single-cell sequencing data was analyzed using the Seurat3 package. For analysis integrating single-cell data from multiple sources, data was imputed using the ALRA package and integrated using mutual nearest-neighbor batch correction. Gene set scores were calculated using the AUCell package and the top quintile of cells were marked as expressing the gene set. Ligand-receptor analysis was performed with the SingleCellSignalR package.

Gene expression signatures were composed of the following. TGFβ(41): *Ccn2, Lox, Spp1, Serpine1, Timp1, Itgb6, Actn1, Thsd1, Fbn1, Mmp14, Tgfbi, Itgav, Mmp3, Tnc, Serpine2, Epha2, Mmp3, Gja4, Rhoa, Axl, Mfap2, Fmod, Marco, Itgb5, Pck1, Ddb2, Hsp90b1, Cd44, Cdc42, Lcp1, Capn2, Ackr3, Ccr2, Cdk4, Tfap2a, Areg, Fscn1, Ctsh, S100a4, Fstl1, Mmp13, Thbs1, Igfbp4, Gadd45a, Scarb1, Tcp1, Hmgb2, Mgp, Gja1, Rfc3*. T-UPR(2): *Spa5, Ddit3, Atf4, Txnip, Bbc3, Pmaip1, Bcl2l11, Bax*. SASP genes(43, 71): *Il7, Cxcl1, Cxcl5, Csf2, Plaur, Timp2, Icam1, Tnfrsf11b, Cxcl2, Ccl2, Igfbp2, Pigf, Tnfrsf1a, Mif, Ccl20, Tnfrsf18, Areg, Serpine1, Plat, Plau, Timp1, Mmp3, Mmp10, Mmp12, Mmp13, Mmp14, Igfbp1, Igfbp3, Igfbp4, Igfbp6, Igfbp7, Fasl, Il6st, Egfr, Il1a, Il1b, Il13, Il15, Cxcl3, Ccl8, Ccl3, Ccl11, Ccl26, Ccl25, Ccl1, Cxcl11, Csf3, Ifng, Cxcl13, Ereg, Nrg1, Egf, Fgf2, Hgf, Fgf7, Vegfa, Ang, Kitl, Cxcl12, Ngf, Ccl7, Clu, Glb1, Lama2, Mmp7, Serpine2, Cdkn2a, Cdkn1a, Tgfb1, Cdkn1c, Cdkn2b, Ccl27a, Lmnb2, Noa1, Tnfrsf10b*.

### Flow cytometry

Lungs were incubated in RPMI supplemented with 80 U/ml DNase I, 2.5 mg/ml collagenase I, 0.4 U/ml dispase, and dissociated in C-tubes on a gentleMACS dissociator (Miltenyi). Cells were stained with: CD45 (30-F11, FITC, Invitrogen 11045182), F4/80 (BM8, PE, Invitrogen 12480182), CD24 (M1/69, PE-Cy7, Invitrogen 25024282), MHC-II IA/IE (M5/114, PerCP-eFluor710, Invitrogen 46532180), CD11b (M1/70, APC-Cy7, Invitrogen A15390), CD11c (N418, APC, Invitrogen 17011482), Ly6C (AL-21, V450, BD 560594), and eFluor506 fixable viability dye (eBioscience 65086614). Data was acquired on a BD FACSverse and analyzed with FlowJo.

### Study approval

The animal model work described in this study was approved by the Institutional Animal Care and Use Committee of the University of California, San Francisco (protocol AN180815).

## Author Contributions

V.C.A., M.T., F.R.P., and D.S. designed the experiments. V.C.A., M.S.D., J.K., T.W., and A.B. conducted the experiments and analyzed the data. V.C.A., F.R.P., and D.S. and wrote the manuscript.

## Acknowledgements

We thank M. Podolsky, R. Ghosh, A. Igbaria, C. Jan, and members of the Papa and Sheppard labs for helpful discussions. We thank the members of the UCSF Genomics CoLab and the UCSF Histology and Biomarker Core for technical support. This work was supported by R01HL145037 (D.S. and F.R.P.), and T32HL007185 and F32HL145990 (V.C.A.).

## Figures

**Figure S1.**
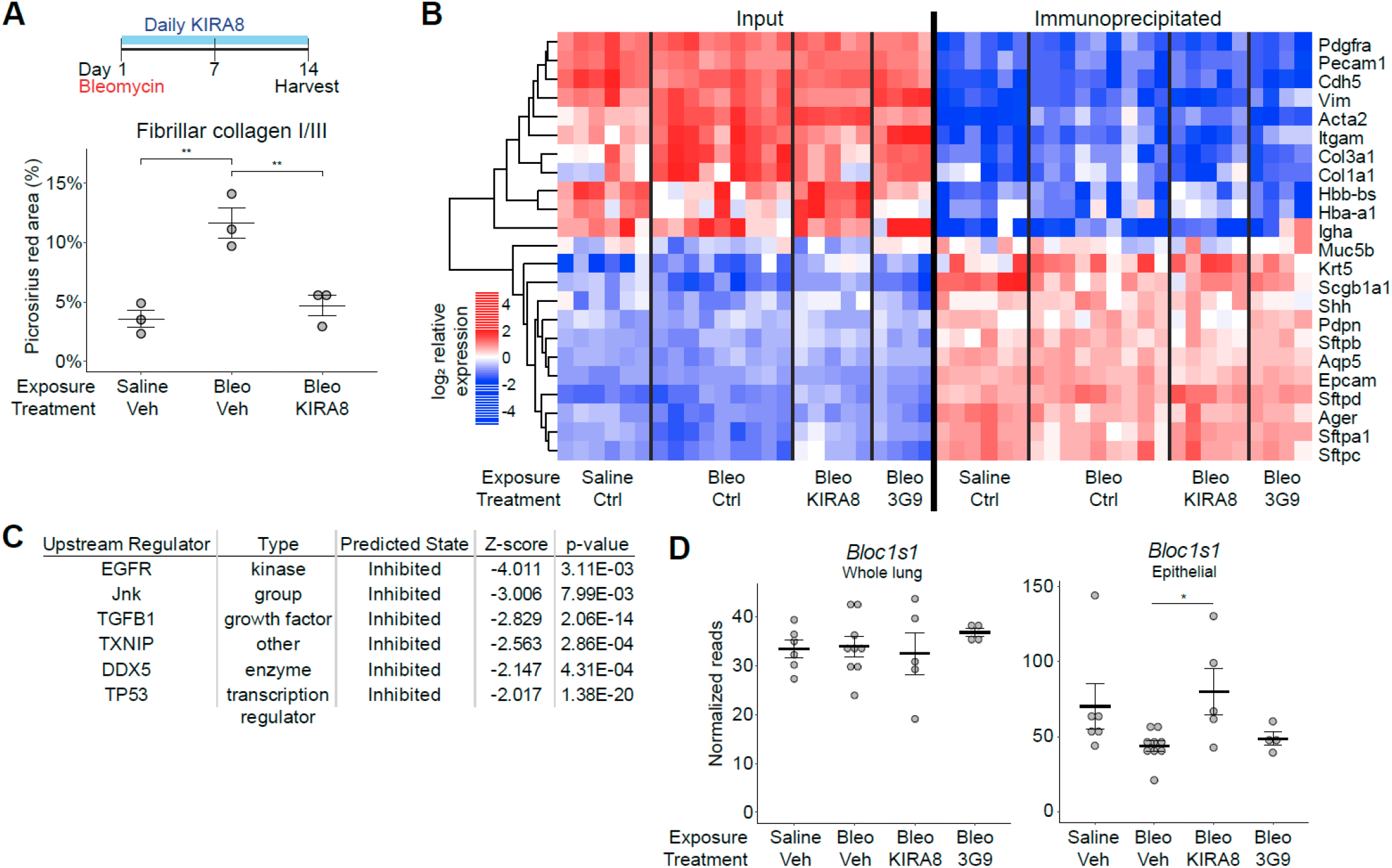
IRE1α regulates fibrosis and profibrotic gene expression in the lung epithelium (related to Figure 1). (A) Histological lung fibrosis after bleomycin exposure and KIRA8 treatment. Mice were exposed to bleomycin, harvested on day 14, and fibrosis measured by picrosirius red staining on formalin-fixed, paraffin-embedded sections. Groups were compared by ANOVA with Fisher’s post-hoc test. (B) Heatmap of cell type specific transcripts in whole lung and epithelial RNAseq libraries. (C) Key Ingenuity Pathway Analysis upstream regulators after KIRA8 treatment in epithelial RiboTag data. (D) *Bloc1s1* (Blos1) expression in whole lung and epithelium after bleomycin exposure and treatment with KIRA8 or 3G9. Reads counts were adjusted for batch and sex. KIRA8 treatment effect was compared by Student’s two-sided t-test. * p<0.05, ** p<0.01.

**Figure S2.**
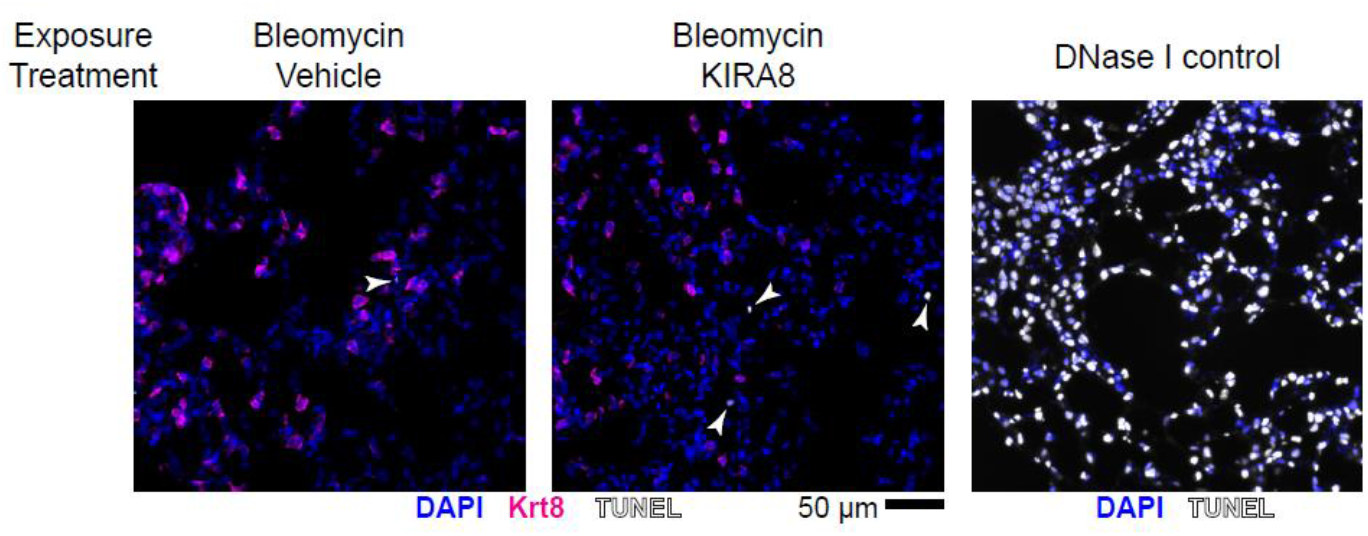
TUNEL staining of frozen lung sections from mice on day 7 after bleomycin exposure and daily treatment with KIRA8. Krt8+ (magenta), TUNEL+ (white) cells are marked by arrowheads (left and center panels). TUNEL staining was validated by DNAse I treatment (right panel).

**Figure S3.**
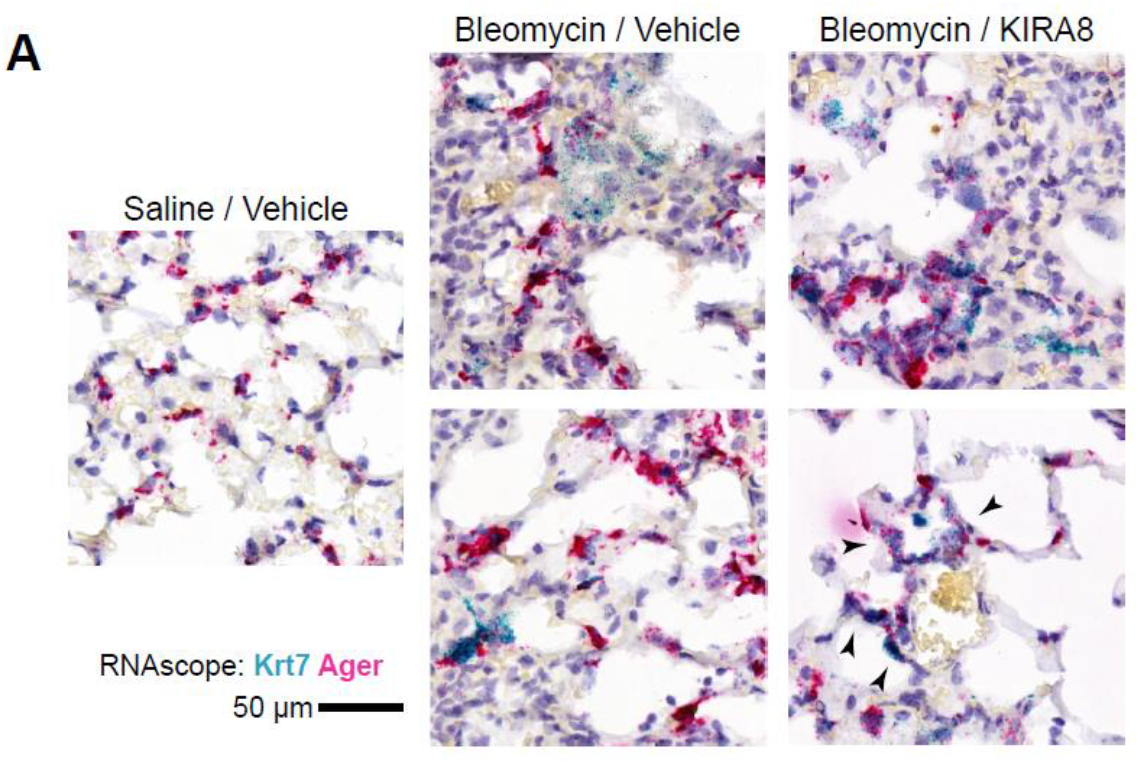
*Krt7+* DATPs with alveolar morphology express the type I alveolar epithelial cell marker *Ager*. RNA in situ hybridization (RNAscope) for *Krt7* (teal) and *Ager* (magenta) at day 14 after bleomycin exposure and treatment with KIRA8, as in Figure 3A. Black arrows mark double-positive cells incorporated into alveolar structures after treatment with bleomycin and KIRA8.

## References

1. Wolters PJ et al. Time for a change: is idiopathic pulmonary fibrosis still idiopathic and only fibrotic? [Internet]. Lancet. Respir. Med. 2018;6(2):154–160.

2. Hetz C, Papa FR. The Unfolded Protein Response and Cell Fate Control. [Internet]. Mol. Cell 2018;69(2):169–181.

3. Han D et al. IRE1α Kinase Activation Modes Control Alternate Endoribonuclease Outputs to Determine Divergent Cell Fates [Internet]. Cell 2009;138(3):562–575.

4. Hollien J et al. Regulated Ire1-dependent decay of messenger RNAs in mammalian cells. [Internet]. J. Cell Biol. 2009;186(3):323–31.

5. Lerner AG et al. IRE1α induces thioredoxin-interacting protein to activate the NLRP3 inflammasome and promote programmed cell death under irremediable ER stress. [Internet]. Cell Metab. 2012;16(2):250–64.

6. Mahavadi P et al. Epithelial stress and apoptosis underlie Hermansky-Pudlak syndrome-associated interstitial pneumonia. [Internet]. Am. J. Respir. Crit. Care Med. 2010;182(2):207–19.

7. Mulugeta S, Nguyen V, Russo SJ, Muniswamy M, Beers MF. A surfactant protein C precursor protein BRICHOS domain mutation causes endoplasmic reticulum stress, proteasome dysfunction, and caspase 3 activation. Am. J. Respir. Cell Mol. Biol. 2005;32(6):521–530.

8. Katzen J et al. A SFTPC BRICHOS mutant links epithelial ER stress and spontaneous lung fibrosis. [Internet]. JCI insight [published online ahead of print: February 5, 2019]; doi:10.1172/jci.insight.126125

9. Borok Z et al. Grp78 loss in epithelial progenitors reveals an age-linked role for endoplasmic reticulum stress in pulmonary fibrosis. Am. J. Respir. Crit. Care Med. 2020;201(2):198–211.

10. Lawson WE et al. Endoplasmic reticulum stress in alveolar epithelial cells is prominent in IPF: association with altered surfactant protein processing and herpesvirus infection. [Internet]. Am. J. Physiol. Lung Cell. Mol. Physiol. 2008;294(6):L1119–26.

11. Korfei M et al. Epithelial Endoplasmic Reticulum Stress and Apoptosis in Sporadic Idiopathic Pulmonary Fibrosis [Internet]. Am. J. Respir. Crit. Care Med. 2008;178(8):838–846.

12. Burman A et al. Localized hypoxia links ER stress to lung fibrosis through induction of C/EBP homologous protein. [Internet]. JCI insight 2018;3(16). doi:10.1172/jci.insight.99543

13. Martinon F, Chen X, Lee A-H, Glimcher LH. TLR activation of the transcription factor XBP1 regulates innate immune responses in macrophages. [Internet]. Nat. Immunol. 2010;11(5):411–8.

14. Qiu Q et al. Toll-like receptor-mediated IRE1α activation as a therapeutic target for inflammatory arthritis. [Internet]. EMBO J. 2013;32(18):2477–90.

15. Blazanin N et al. ER stress and distinct outputs of the IRE1α RNase control proliferation and senescence in response to oncogenic Ras. Proc. Natl. Acad. Sci. U. S. A. 2017;114(37):9900–9905.

16. Dufey E et al. Genotoxic stress triggers the activation of IRE1α-dependent RNA decay to modulate the DNA damage response [Internet]. Nat. Commun. 2020;11(1):2401.

17. Tam AB, Koong AC, Niwa M. Ire1 has distinct catalytic mechanisms for XBP1/HAC1 splicing and RIDD [Internet]. Cell Rep. 2014;9(3):850–858.

18. Chen X et al. XBP1 promotes triple-negative breast cancer by controlling the HIF1α pathway. [Internet]. Nature 2014;508(7494):103–107.

19. Pluquet O et al. Posttranscriptional regulation of per1 underlies the oncogenic function of IREα. Cancer Res. 2013;73(15):4732–4743.

20. Heindryckx F et al. Endoplasmic reticulum stress enhances fibrosis through IRE1α-mediated degradation of miR-150 and XBP-1 splicing [Internet]. EMBO Mol. Med. 2016;8(7):729–744.

21. Gonzalez-Teuber V et al. Small Molecules to Improve ER Proteostasis in Disease. [Internet]. Trends Pharmacol. Sci. 2019;40(9):684–695.

22. Thamsen M et al. Small molecule inhibition of IRE1α kinase/RNase has anti-fibrotic effects in the lung. [Internet]. PLoS One 2019;14(1):e0209824.

23. Harrington PE et al. Unfolded Protein Response in Cancer: IRE1α Inhibition by Selective Kinase Ligands Does Not Impair Tumor Cell Viability. [Internet]. ACS Med. Chem. Lett. 2015;6(1):68–72.

24. Morita S et al. Erratum: Targeting ABL-IRE1α Signaling Spares ER-Stressed Pancreatic β Cells to Reverse Autoimmune Diabetes (Cell Metabolism (2017) 25(4) (883–897.e8) (S1550413117301705) (10.1016/j.cmet.2017.03.018)) [Internet]. Cell Metab. 2017;25(5):1207.

25. Sanz E et al. Cell-type-specific isolation of ribosome-associated mRNA from complex tissues [Internet]. Proc. Natl. Acad. Sci. 2009;106(33):13939–13944.

26. Haimon Z et al. Re-evaluating microglia expression profiles using RiboTag and cell isolation strategies. [Internet]. Nat. Immunol. 2018;19(6):636–644.

27. Weinreb PH et al. Function-blocking Integrin α v β 6 Monoclonal Antibodies [Internet]. J. Biol. Chem. 2004;279(17):17875–17887.

28. Urano F et al. Coupling of stress in the ER to activation of JNK protein kinases by transmembrane protein kinase IRE1. [Internet]. Science 2000;287(5453):664–6.

29. Moore K, Hollien J. Ire1-mediated decay in mammalian cells relies on mRNA sequence, structure, and translational status. Mol. Biol. Cell 2015;26(16):2873–2884.

30. Upton J-P et al. IRE1α cleaves select microRNAs during ER stress to derepress translation of proapoptotic Caspase-2. [Internet]. Science 2012;338(6108):818–22.

31. Lu Y, Thomson JM, Wong HYF, Hammond SM, Hogan BLM. Transgenic over-expression of the microRNA miR-17-92 cluster promotes proliferation and inhibits differentiation of lung epithelial progenitor cells. Dev. Biol. 2007;310(2):442–453.

32. Li Y, Choi PS, Casey SC, Dill DL, Felsher DW. MYC through miR-17-92 suppresses specific target genes to maintain survival, autonomous proliferation, and a Neoplastic state [Internet]. Cancer Cell 2014;26(2):262–272.

33. Du WW et al. MiR-17 extends mouse lifespan by inhibiting senescence signaling mediated by MKP7. Cell Death Dis. 2014;5(7):1–14.

34. Faraonio R et al. A set of miRNAs participates in the cellular senescence program in human diploid fibroblasts [Internet]. Cell Death Differ. 2012;19(4):713–721.

35. Li G, Luna C, Qiu J, Epstein DL, Gonzalez P. Alterations in microRNA expression in stress-induced cellular senescence [Internet]. Mech. Ageing Dev. 2009;130(11–12):731–741.

36. Mestdagh P et al. The miR-17-92 MicroRNA Cluster Regulates Multiple Components of the TGF-β Pathway in Neuroblastoma. Mol. Cell 2010;40(5):762–773.

37. Agarwal V, Bell GW, Nam J-W, Bartel DP. Predicting effective microRNA target sites in mammalian mRNAs. [Internet]. Elife 2015;4(AUGUST 2015):1–38.

38. Baek HA et al. Involvement of endoplasmic reticulum stress in myofibroblastic differentiation of lung fibroblasts. [Internet]. Am. J. Respir. Cell Mol. Biol. 2012;46(6):731–9.

39. Kim RS et al. The XBP1 Arm of the Unfolded Protein Response Induces Fibrogenic Activity in Hepatic Stellate Cells Through Autophagy. [Internet]. Sci. Rep. 2016;6(December):39342.

40. Chen Y-T et al. Endoplasmic reticulum protein TXNDC5 promotes renal fibrosis by enforcing TGFβ signaling in kidney fibroblasts. J. Clin. Invest. [published online ahead of print: 2021]; doi:10.1172/jci143645

41. Kaminski N et al. Global analysis of gene expression in pulmonary fibrosis reveals distinct programs regulating lung inflammation and fibrosis. [Internet]. Proc. Natl. Acad. Sci. U. S. A. 2000;97(4):1778–83.

42. Coppé J-P et al. Senescence-Associated Secretory Phenotypes Reveal Cell-Nonautonomous Functions of Oncogenic RAS and the p53 Tumor Suppressor [Internet]. PLoS Biol. 2008;6(12):e301.

43. Yao C et al. Senescence of alveolar stem cells drives progressive pulmonary fibrosis. bioRxiv 2019;

44. Lehmann M et al. Senolytic drugs target alveolar epithelial cell function and attenuate experimental lung fibrosis ex vivo [Internet]. Eur. Respir. J. 2017;50(2). doi:10.1183/13993003.02367-2016

45. Strunz M et al. Alveolar regeneration through a Krt8+ transitional stem cell state that persists in human lung fibrosis [Internet]. Nat. Commun. 2020;11(1). doi:10.1038/s41467-020-17358-3

46. Kathiriya JJ, Brumwell AN, Jackson JR, Tang X, Chapman HA. Distinct Airway Epithelial Stem Cells Hide among Club Cells but Mobilize to Promote Alveolar Regeneration [Internet]. Cell Stem Cell 2020;26(3):346–358.e4.

47. Kobayashi Y et al. Persistence of a regeneration-associated, transitional alveolar epithelial cell state in pulmonary fibrosis [Internet]. Nat. Cell Biol. 2020;22(8):934–946.

48. Jiang P et al. Ineffectual Type 2-to-Type 1 Alveolar Epithelial Cell Differentiation in Idiopathic Pulmonary Fibrosis: Persistence of the KRT8hi Transitional State. Am. J. Respir. Crit. Care Med. 2020;201(11):1443–1447.

49. Choi J et al. Inflammatory Signals Induce AT2 Cell-Derived Damage-Associated Transient Progenitors that Mediate Alveolar Regeneration [Internet]. Cell Stem Cell 2020;27(3):366–382.e7.

50. Schafer MJ et al. Cellular senescence mediates fibrotic pulmonary disease [Internet]. Nat. Commun. 2017;8(1):14532.

51. Gibbons MA et al. Ly6Chi monocytes direct alternatively activated profibrotic macrophage regulation of lung fibrosis. Am. J. Respir. Crit. Care Med. 2011;184(5):569–581.

52. Misharin A V. et al. Monocyte-derived alveolar macrophages drive lung fibrosis and persist in the lung over the life span [Internet]. J. Exp. Med. 2017;214(8):2387–2404.

53. Aran D et al. Reference-based analysis of lung single-cell sequencing reveals a transitional profibrotic macrophage [Internet]. Nat. Immunol. 2019;20(2):163–172.

54. Satoh T et al. Identification of an atypical monocyte and committed progenitor involved in fibrosis. Nature 2017;541(7635):96–101.

55. Misharin A V. et al. Monocyte-derived alveolar macrophages drive lung fibrosis and persist in the lung over the life span. J. Exp. Med. 2017;214(8):2387–2404.

56. Habermann AC et al. Single-cell RNA-sequencing reveals profibrotic roles of distinct epithelial and mesenchymal lineages in pulmonary fibrosis [Internet]. bioRxiv 2019;753806.

57. Habermann A et al. Single-cell RNA sequencing reveals profibrotic roles of distinct epithelial and mesenchymal lineages in pulmonary fibrosis. Sci. Adv. 2020;6(28):eaba1972.

58. Lawson WE et al. Endoplasmic reticulum stress enhances fibrotic remodeling in the lungs. [Internet]. Proc. Natl. Acad. Sci. U. S. A. 2011;108(26):10562–7.

59. Auyeung VC, Sheppard D. Stuck in a Moment: Does Abnormal Persistence of Epithelial Progenitors Drive Pulmonary Fibrosis? [Internet]. Am. J. Respir. Crit. Care Med. 2020;rccm.202010-3898ED.

60. Verheyden JM, Sun X. A transitional stem cell state in the lung. [Internet]. Nat. Cell Biol. 2020;22(9):1025–1026.

61. Romero-Ramirez L et al. XBP1 is essential for survival under hypoxic conditions and is required for tumor growth. Cancer Res. 2004;64(17):5943–5947.

62. Anderson LL, Mao X, Scott B a, Crowder CM. Survival from Hypoxia in C. elegans by Inactivation of Aminoacyl-tRNA Synthetases [Internet]. Science (80-.). 2009;323(5914):630–633.

63. Richardson CE, Kooistra T, Kim DH. An essential role for XBP-1 in host protection against immune activation in C. elegans. Nature 2010;463(7284):1092–1095.

64. Simic MS et al. Transient activation of the UPRER is an essential step in the acquisition of pluripotency during reprogramming. Sci. Adv. 2019;5(4). doi:10.1126/sciadv.aaw0025_rfseq1

65. Dews M et al. The Myc-miR-17∼92 axis blunts TGFβ signaling and production of multiple TGFβ-dependent antiangiogenic factors. Cancer Res. 2010;70(20):8233–8246.

66. Ernst A et al. De-repression of CTGF via the miR-17-92 cluster upon differentiation of human glioblastoma spheroid cultures. [Internet]. Oncogene 2010;29(23):3411–22.

67. Ivanovska I et al. MicroRNAs in the miR-106b Family Regulate p21/CDKN1A and Promote Cell Cycle Progression. Mol. Cell. Biol. 2008;28(7):2167–2174.

68. Ventura A et al. Targeted Deletion Reveals Essential and Overlapping Functions of the miR-17∼92 Family of miRNA Clusters. Cell 2008;132(5):875–886.

69. Dakhlallah D et al. Epigenetic Regulation of miR-17∼92 Contributes to the Pathogenesis of Pulmonary Fibrosis [Internet]. Am. J. Respir. Crit. Care Med. 2013;187(4):397–405.

70. Vaughan A, Vaughan A, Brumwell A, Chapman H. Lung Epithelial Cell Prep [Internet]. Protoc. Exch. [published online ahead of print: February 23, 2015]; doi:10.1038/protex.2015.017

71. Coppé J-P, Desprez P-Y, Krtolica A, Campisi J. The Senescence-Associated Secretory Phenotype: The Dark Side of Tumor Suppression. Annu. Rev. Pathol. Mech. Dis. 2010;5(1):99–118.

72. Misharin A V., Morales-Nebreda L, Mutlu GM, Budinger GRS, Perlman H. Flow cytometric analysis of macrophages and dendritic cell subsets in the mouse lung. Am. J. Respir. Cell Mol. Biol. 2013;49(4):503–510.

